# Deep generative design of neutralizing nanobodies against SARS-CoV-2 variants

**DOI:** 10.1101/2024.10.29.620982

**Authors:** Liyun Huo, Tian Tian, Yanqin Xu, Qin Qin, Xinyi Jiang, Qiang Huang

## Abstract

In recent years, single-domain heavy chain antibodies (nanobodies), with only one-tenth the molecular weight of conventional antibodies, have emerged as important therapeutic proteins in the fight against SARS-CoV-2. However, the rapid mutation of the virus often renders existing nanobodies ineffective, underscoring the need to develop nanobodies that specifically target new variants. Traditional methods for discovering nanobodies are time-consuming and complex, making it difficult to efficiently identify nanobodies that bind to specific epitopes. To address this, we propose a de novo nanobody design method based on a Generative Adversarial Network (GAN). We developed a deep generative model, AiCDR, which consists of three discriminators and one generator. By determining the contribution of these three discriminators to the entire network, we can enhance the discrimination of generated sequences and reduce the similarity of the generated CDR3 sequences with natural peptides and random sequences, thereby ensuring their nature-like properties. These generated CDR3 sequences were then grafted onto humanized nanobody scaffolds, resulting in a structural library of approximately 10^4^ nanobodies with natural-like properties. Using computational methods, we screened this library against the Spike (S) protein of the SARS-CoV-2 Omicron variant and identified 10 candidate nanobodies. Functional assays confirmed that two of these nanobodies exhibited neutralizing activity against the S protein. Our study demonstrates the potential of deep learning in nanobody design and offers a novel approach for developing nanobodies that target rapidly evolving viral variants.

## Introduction

In recent years, the global pandemic of SARS-CoV-2 has led to a severe COVID-19 crisis, significantly impacting public health and the global economy. Researchers have accelerated vaccine development and pursued various antiviral treatments targeting the virus. (*1*) Among these treatments, neutralizing antibodies are particularly essential, effectively inhibiting viral transmission by blocking the binding of the viral Spike (S) protein to the ACE2 receptor on human host cells. (*2*) To date, the FDA has approved eight neutralizing antibodies for COVID-19 treatment. Additionally, single-domain heavy chain antibodies derived from camelid species, also known as nanobodies, (*3*) have gained attention due to their unique structural characteristics. Numerous studies have demonstrated that nanobodies can efficiently inhibit S protein–ACE2 binding, thus impeding viral spread. (*4*) With a molecular weight approximately one-tenth that of traditional antibodies (around 15 kDa), nanobodies exhibit enhanced tissue penetration, low immunogenicity, and ease of large-scale production. (*5, 6*) Due to their compact structure, nanobodies are capable of binding antigenic epitopes that are often inaccessible to conventional antibodies, particularly conserved epitopes, making them advantageous for SARS-CoV-2 neutralization.

However, the rapid evolution of SARS-CoV-2 presents challenges to the efficacy of existing nanobodies. Since its discovery, SARS-CoV-2 has developed multiple variants, including Alpha, Beta, Gamma, Delta, and Omicron. (*7*) Among these, Omicron stands out for its high transmissibility and immune escape capabilities. The initial Omicron subvariant BA.1 contains 35 mutations on the Spike protein, significantly enhancing immune evasion and transmissibility. (*8*) With ongoing evolution, Omicron now includes over 300 subvariants, including XBB, which exhibit resistance to all currently available monoclonal antibodies. (*9*) Consequently, developing novel nanobodies that can target these continually evolving variants is essential.

Traditional methods for nanobody discovery typically involve library construction and experimental screening. Library construction generally includes immunized, naïve, and synthetic libraries. Immunized libraries require antigen injections in camelid species to generate antibodies against specific epitopes, (*5*) while naïve libraries necessitate a large number of animals and fresh blood samples to construct sufficiently large nanobody libraries. (*10*) Synthetic libraries rely on artificial sequence synthesis to mimic natural antibody features, yet they often exhibit weaker physicochemical properties and require extended development periods. Although mature experimental screening techniques such as phage display and yeast display are available, they remain costly, time-consuming, and labor-intensive, with limited capacity to produce antibodies targeting specific epitopes. As a result, developing more efficient and precise antibody screening methods has become a key research focus.

Recent computational approaches have introduced new perspectives in antibody development. Synthetic libraries typically rely on extracted features from natural sequences to generate artificial sequences, thus eliminating the need for animal immunization and reducing both cost and development time. (*11, 12*) However, synthetic sequences are still different from natural sequences in terms of physicochemical properties. (*10*) Generative models like Generative Adversarial Networks (GANs) (*13*) have demonstrated significant potential in antibody design, with success in the generation of peptide and protein sequences. (*14, 15*) Previous studies have shown that CDR3 grafting can achieve affinity transfer, (*16*) and recent efforts have demonstrated that de novo design of antibody CDR3 sequences can enable binding to target proteins. (*17*) These findings underscore the promising future of generating natural-like CDR3 sequences for target protein binding.

In this study, we propose a GAN-based method for de novo design of nanobody CDR3 sequences. We developed a generative model, AiCDR, capable of producing natural-like CDR3 sequences, which we grafted onto a humanized antibody scaffold to construct a library of approximately 10^4^ nanobody structures. Through computational screening, we identified nanobody candidates capable of binding to the SARS-CoV-2 Omicron BA.1 S protein, ultimately isolating ten candidates. Functional assays confirmed that two of these nanobodies exhibited neutralization activity against the Omicron S protein.

## Results

### Network architecture of AiCDR

To generate natural-like CDR3 sequences, we constructed a GAN model, AiCDR, based on the hierarchical reinforcement learning framework of LeakGAN. (*18*) As shown in the top of Fig. 1, the LeakGAN network consists of a Discriminator and a Generator. Different from the valina GAN, LeakGAN’s Discriminator continuously leaks features of the training samples to the Generator during the training process, guiding the Generator to produce highquality sequences. While the LeakGAN network was designed to work effectively with positive samples, its ability to distinguish negative samples could be further enhanced. To improve this capability, we added external modules inspired by PepGAN, (*14*) and then make the entire network more powerful in distinguishing positive and negative sequences by confirming their contribution to the overall network.

**Figure 1.**
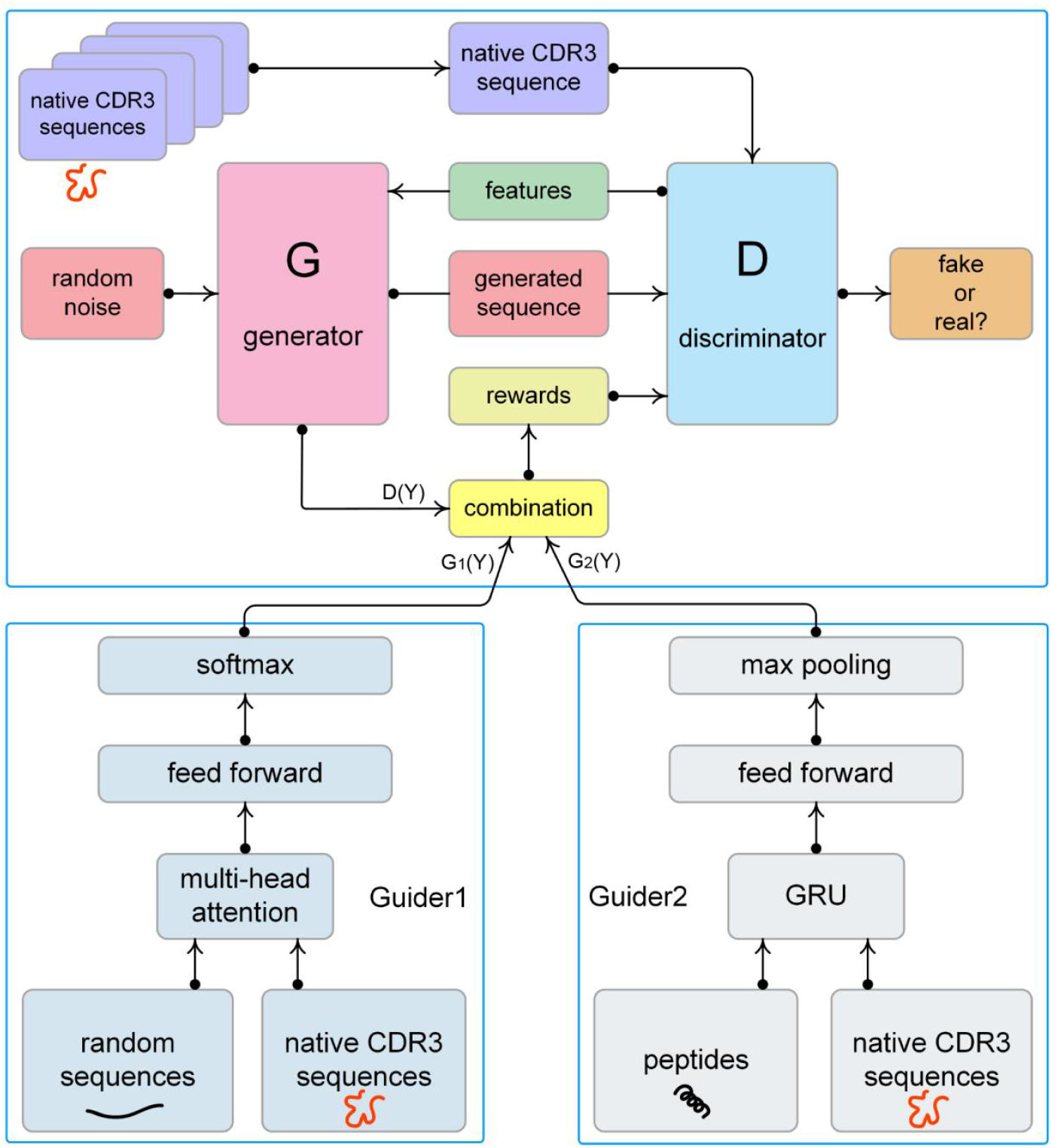
Neural network architecture of the AiCDR model. The network consists of four main components: Discriminator, Generator, Guider1, and Guider2. The Discriminator (blue) is trained on native CDR3 sequences and provides its recognized features of native CDR3 to the Generator. Guider1 (light green) and Guider2 (light blue) are auxiliary modules of the Discriminator. Guider1 is constructed based on the multi-attention mechanism, which is trained with random sequences against native CDR3 sequences and is used to discriminate between random sequences and CDR3 sequences, while Guider2 is constructed by the GRU network and is trained with peptide sequences against native CDR3 sequences to discriminate between individual peptide sequences and native CDR3 sequences that are part of nanobodies. These two modules can further enhance the Discriminator to determine if the input sequence is a native CDR3. Discriminator, Guider1 and Guider2 calculate rewards based on different ratios to guide the generator (purple) to generate natural-like CDR3 sequences.

To incorporate negative samples into the training process, we added two external modules to AiCDR. As shown in Fig. 1 (bottom), these modules, named Guider1 and Guider2, were designed to improve the network’s ability to distinguish between different sequence types. Guider1 consists of a multi-head self-attention mechanism with 8 heads and two fully connected layers, while Guider2 is a Gated Recurrent Unit (GRU) with 256 neurons. We trained Guider1 using random sequences as negative samples. Given that CDR3 is a fragment of the nanobody, with a length similar to full-length peptides, we used full-length peptide sequences as negative samples to train Guider2. Both Guider modules were trained with natural CDR3 sequences as positive samples. This setup allowed AiCDR to distinguish between random and functional sequences, as well as between fragmented and full-length sequences. The Guider modules contribute to the calculation of the reward function, which consists of three key components in this model:

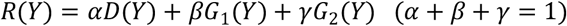

where Y is the sequence generated by Generator, D(Y) is the output of Discriminator network for evaluating the authenticity of the sequence Y; G1(Y) and G2(Y) are the outputs of Guider1, and Guider2, respectively, both are the probability that Y is a natural CDR3 sequence.

### Training process of AiCDR

To construct the training datasets, we prepared three sets for model training. First, we collected 2,526,950 nanobody sequences from the INDI (*19*) database as of April 25, 2023. Since the CDR3 sequence lengths predominantly range between 5 ∼25 a.a., (*19*) we retained only those sequences within this range and used CD-HIT (*20*) to eliminate redundant sequences with more than 90% similarity, resulting in 1,702,270 unique CDR3 sequences. Next, we collected 1,839,619 peptide sequences from the NCBI (*21*) database, again retaining only those within the 5 ∼ 25 a.a. range, removing high-similarity sequences, and leaving 107,562 peptide sequences. Finally, we randomly generated 1,702,270 sequences between 5 ∼ 25 a.a. to serve as random sequences.

To train the Guider module, we use 1,702,270 CDR3 sequences and 1,702,270 random sequences as the training set for the Guider1 network; 107,562 CDR3 sequences and 107,562 polypeptide sequences are randomly selected as the training set for the Guider2 network (Table 1). We train the two Guider modules separately. When Guider1 is trained to 10,000 times, the loss and accuracy of the training set and the validation set are basically unchanged, and the network reaches equilibrium (Fig. S1a-b); while when Guider2 is trained to 52 times, the loss of the training set and the validation set reaches the minimum (loss = 0.3), and the accuracy reaches the maximum (accuracy = 0.9). At this time, we stop the training of this model (Fig. S1c-d).

**Table 1.** Training dataset for the AiCDR model.

| Module | Sequence type | Total number |
| --- | --- | --- |
| Guider1 |  |  |
| Positive | Native CDR3 | 1,702,270 |
| Negative | Random | 1,702,270 |
| Guider2 |  |  |
| Positive | Native CDR3 | 107,562 |
| Negative | Individual peptide | 107,562 |
| Generator & Discriminator | Native CDR3 | 1,702,270 |

To determine optimal values for the parameters α, β, and γ in the reward function, defining the contributions of Guider1, Guider2, and the Discriminator in evaluating generated sequences, we conducted the following experiments. Initially, we used 1,702,270 CDR3 sequences as training data and established 61 combinations of α, β, and γ values with α values ranging from 0 to 1 in increments of 0.1(Table S1), training each network configuration for 200 epochs. The quality of generated sequences was assessed based on their similarity to those in the training set, enabling us to determine whether the model training was complete. Specifically, at each epoch, AiCDR generated 9,984 sequences, which were subsequently compared with the 1,702,270 training sequences using BLAST (E-value = 1e -5). The resulting number of similar sequences for each epoch was plotted as a line graph (Fig. S2a). As shown in the figure, the combination AiCDR (α = 0.6, β = 0.3, γ = 0.1), abbreviated as 631 in the figure, produced the highest number of similar sequences at the 198th epoch, reaching a total of 284. Notably, when α, β, and γ were set to zero, AiCDR functioned as a basic LeakGAN network with a single discriminator, which generated fewer similar sequences than other AiCDR models. Based on this analysis, we determined the optimal values for the three parameters to be α = 0.6, β = 0.3, and γ = 0.1.

With the hyperparameter values of the reward function confirmed, we continued training AiCDR (α = 0.6, β = 0.3, γ = 0.1) for an additional 100 epochs, reaching a total of 300 epochs. As shown in Fig. S2b, AiCDR generated 284 similar sequences at the 198th epoch and 213 at the 261st epoch. From the 200th to the 258th epochs and from the 265th to the 300th epochs, fewer than 50 similar sequences were generated. These results across 300 epochs indicate that the highest number of similar sequences, 284, was achieved at the 198th epoch, marking this as the completion point for model training.

### Analysis of Naturalness and Diversity in generated CDR3 Sequence

To assess the quality of CDR3 sequences generated by the AiCDR model, we analyze the naturalness and diversity of these sequences. For naturalness, we performed statistical analyses in four areas: sequence length, amino acid distribution, proportion of paratope hotspot amino acids, and nine physicochemical properties. First, we analyzed the length of the 9,984 generated sequences (Fig. 2a). As shown, all generated sequences ranged from 5 to 25 amino acids, consistent with the training dataset. The length distribution of the generated sequences (red) closely aligned with that of the training set (blue), both peaking between 16 and 20 amino acids and exhibiting the fewest sequences between 5 and 13 or 23 and 25 amino acids. Next, we evaluated the amino acid composition of the generated sequences in comparison to that of the training set. The amino acid frequency distribution in Fig. 2b reveals that both generated and training sets exhibited similar compositions. Additionally, we examined the count of paratope hotspot amino acids: alanine (A), serine (S), threonine (T), tyrosine (Y), and tryptophan (W) in both generated and training sequences, with box plots illustrating their proportion in each set (Fig. 2c). The distributions were analogous, with alanine (A) being the most abundant, followed by serine (S) and threonine (T). Finally, we computed nine physicochemical properties (including length, molecular weight, aromaticity, stability, isoelectric point, charge, and secondary structure preferences) for both generated and natural sequences, visualized using t-distributed stochastic neighbor embedding (t-SNE) (Fig. 2d). The results indicate that the generated CDR3s possess properties comparable to those of natural CDR3s. These findings suggest that the generated CDR3 sequences share similar length distributions, amino acid composition, and physicochemical attributes with natural CDR3 sequences. Moreover, the proportion of hotspot amino acids in the generated CDR3 sequences, essential for antigen binding in nanobodies, closely matched that in natural sequences, suggesting high naturalness in AiCDR-generated sequences.

**Figure 2.**
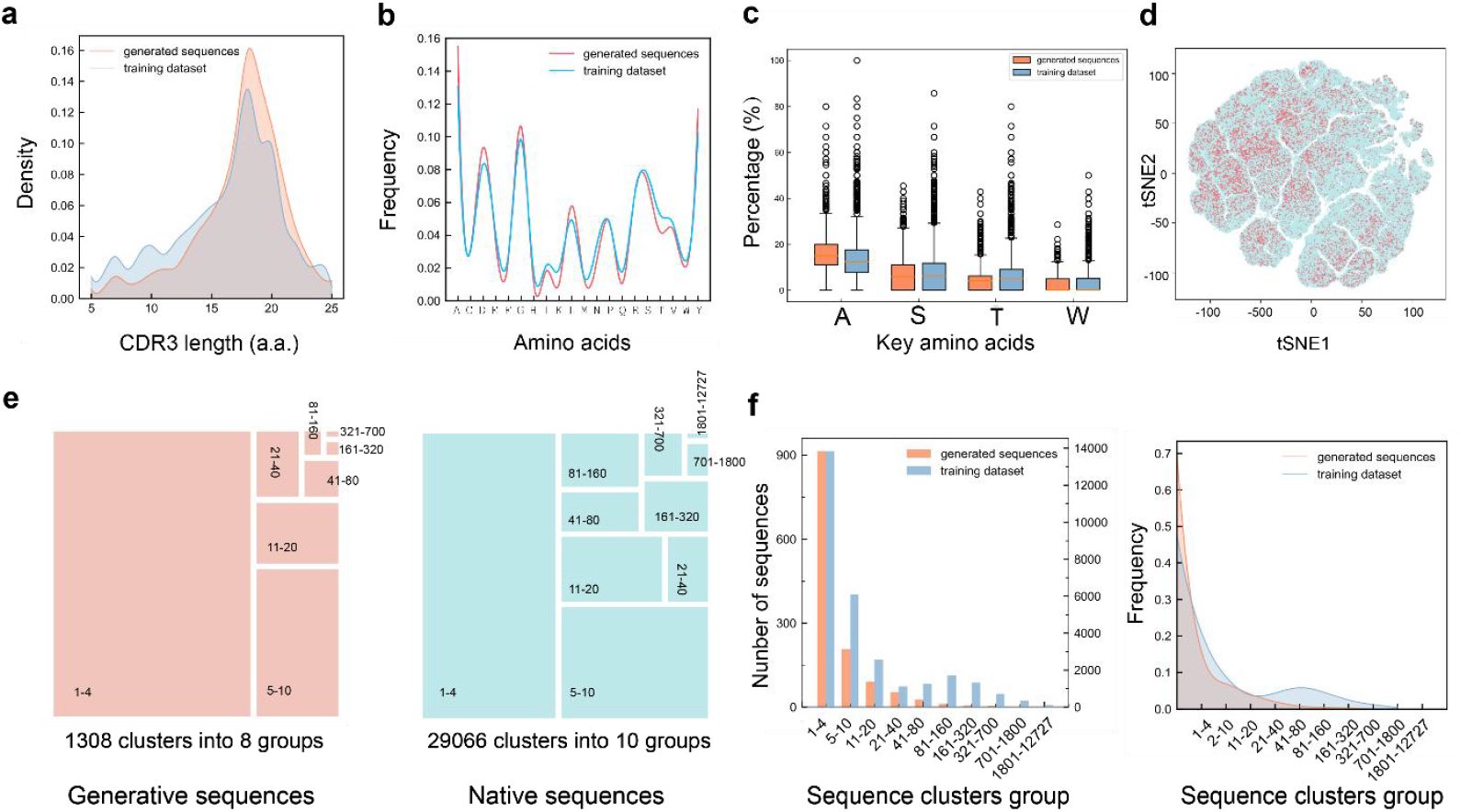
Naturalness and diversity analyses of sequences generated from scratch. (a) CDR3 length distribution. The length distribution of CDR3 sequences generated by AiCDR (red) is comparable to that of the training dataset (blue). All of generated sequences have lengths centred on 5-25 amino acids, which is consistent with the training dataset. (b) Amino acid composition of sequences. Generated sequences show a highly similar amino acid frequency and compositional variability to the training dataset. (c) Hot amino acid statistics. The ratio of A, S, T and W amino acids in the generated sequence (purple) and the training set sequence (green) is almost identical. (d) t-SNE analysis of physiochemical features of sequences (perplexity=100). The nine major physiochemical features (length, molecular weight, aromaticity, stability, isoelectric point, charge, Tendency of secondary structures to helixes, turns or sheets) were similarly distributed between the two types of sequences. (e) Diversity analysis: 1,308 clusters of generated sequences were divided into eight major groups, while 29,066 clusters of native sequences were divided into ten major groups. (f) The number and proportion of clusters for the generated sequence (orange) and the training set sequence (green).

To evaluate sequence diversity, we performed clustering analysis on the 9,984 generated CDR3 sequences and the 1,702,270 natural CDR3 sequences from the training set using CD-HIT (identity = 0.5) (Table S2). The generated sequences formed 1,308 clusters, and the natural sequences formed 29,066 clusters. We then categorized these clusters based on their sequence count, with clusters containing 1 ∼ 4 sequences grouped together, clusters with 5 ∼ 10 sequences were grouped into a separate category, and so on (Table S2). Fig. 2e illustrates that the generated sequences were divided into eight groups (red), with the largest proportion comprising clusters of 1 ∼ 4 sequences, followed by clusters of 5 ∼ 10 sequences. Similarly, the natural sequences (blue) also divided into different groups, with the largest proportion in clusters containing 1 ∼ 4 sequences, followed by 5 ∼ 10 sequence clusters. The larger dataset of natural sequences also included additional groups with clusters containing 700 ∼ 1,800 and 1,800 ∼12,727 sequences. To describe the proportions occupied by these groups quantitatively, we calculated the number and percentage of each group. Fig. 2f shows that approximately 68% (3,500) of the generated sequences formed 900 clusters, each containing 1 ∼ 4 sequences. The number of clusters within each group decreased as the sequence count per cluster increased. For the training set, around 48% (54,000) of natural sequences formed 14,000 clusters with 1 ∼ 4 sequences, also showing fewer clusters as sequence count per cluster increased. These analyses indicate that AiCDR-generated sequences exhibit diversity levels comparable to those of natural sequences.

### Construction of a Nanobody Structural Library with Diverse CDR3 Sequences

Based on the previous analysis, we observed that sequences generated by AiCDR possess both high naturalness and substantial diversity. Consequently, we leveraged these generated CDR3 sequences to construct a nanobody structural library (Fig. 3). Initially, we selected Caplacizumab (*22*) as the framework for CDR3 sequence grafting, as it is a nanobody therapeutic for acquired thrombotic thrombocytopenic purpura (aTTP). Utilizing Caplacizumab as a grafting scaffold reduces immunogenicity risks, thereby facilitating later stages of drug development. Additionally, in our laboratory, we achieved soluble expression of Caplacizumab, and subsequent Biolayer Interferometry (BLI) analysis confirmed its lack of binding affinity for the Omicron RBD (Fig. S7a). This validation supports Caplacizumab’s use as a scaffold to assess whether the CDR3 sequences generated can indeed mediate nanobody binding to the Omicron RBD.

**Figure 3.**
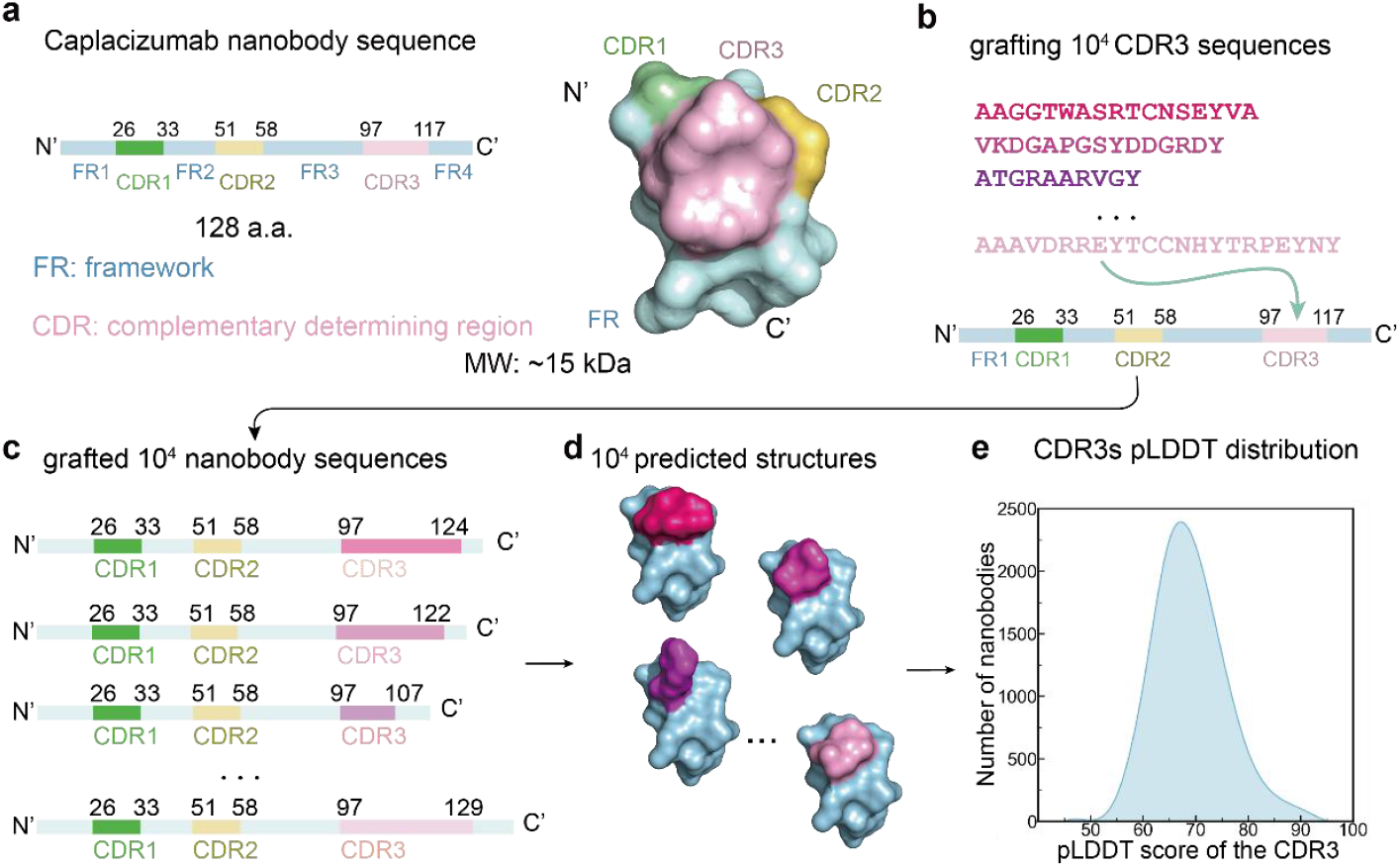
Design of CDR3 grafting for full-length nanobody. (a) Sequence and corresponding nanobody structure of Caplacizumab. (b) Process of grafting nanobody CDR3 sequences. the 9984 CDR3 sequences generated by AiCDR were grafted individually onto the CDR3 region of Caplacizumab. (c) 9984 sequences obtained after grafting. (d) Structural library of 104 nanobodies obtained using structure prediction. (e) Distribution of pLDDT scores for predicted CDR3 structures. pLDDT scores were calculated to assess the structural reliability.

Caplacizumab comprises 128 amino acids with an approximate molecular weight of 15 kDa. We designated Caplacizumab’s sequence [Protein Data Bank (PDB) 7EOW] into framework and CDR regions by IMGT (*23*) numbering, identifying residues 97 to 117 as the CDR3 domain (Fig. 3a). Then, We grafted each of the 9,984 generated CDR3 sequences individually onto Caplacizumab’s CDR3 domain (97 ∼ 117 residues; Fig. 3b), producing 9,984 unique nanobody sequences distinguished only by their CDR3 sequences (Fig. 3c). Using Alphafold2, (*24*) we subsequently predicted the structures of these nanobodies, yielding a structural library of approximately (10^4^) nanobody configurations (Fig. 3d).

As illustrated in Fig. S3, the framework regions within these nanobodies exhibited consistently high pLDDT values, exceeding 95, indicating robust structural reliability in the framework. Conversely, the pLDDT values for the three CDR domains were comparatively lower, with the lowest values observed in the CDR3 region. Therefore, we calculated the average pLDDT values specifically for the CDR3 regions. As depicted in Fig. 3e, approximately 2,400 nanobodies demonstrated an average CDR3 pLDDT of 68, with all generated CDR3 regions displaying average pLDDT values between 45 and 94. This outcome aligns with existing literature, (*25*) as such lower pLDDT values are expected for flexible regions like CDR3. Accordingly, we employed this structural library of 10^4^ nanobodies for the next phase of computational screening.

### Nanobody Screening Targeting SARS-CoV-2 Omicron S protein

To identify nanobodies in the constructed structural library that target the SARS-CoV-2 Omicron Spike (S) protein, we conducted computational screening, as illustrated in Fig. 4a. To enhance the efficiency of the screening process, we selected the receptor-binding domain (RBD) (PDB 7QTK) of the Omicron BA.1 S protein, which interacts with the human ACE2 receptor, as the docking target. Because the S protein forms a trimer, using a monomeric RBD in computational screening requires consideration of binding sites and spatial constraints between monomers to improve the accuracy of the results. Since the probability of the Omicron BA.1 RBD existing in the “up” state reaches 95%, (*26*) exposing a greater surface area for potential nanobody interaction, we chose the “up” RBD within the S protein as the target protein. To avoid steric hindrance when nanobodies bind the S protein, we identified amino acids within 4 Å of the RBD that interface with the two adjacent “down” RBDs and the N-terminal domain (NTD) (*27*) (Fig. S4). By excluding these residues, we defined the remaining amino acids on the RBD surface as the target epitope, primarily encompassing the following regions: 132 ∼ 155, 237 ∼ 254, and 275 ∼ 294 a.a. (Fig. 4a, red-highlighted areas).

**Figure 4.**
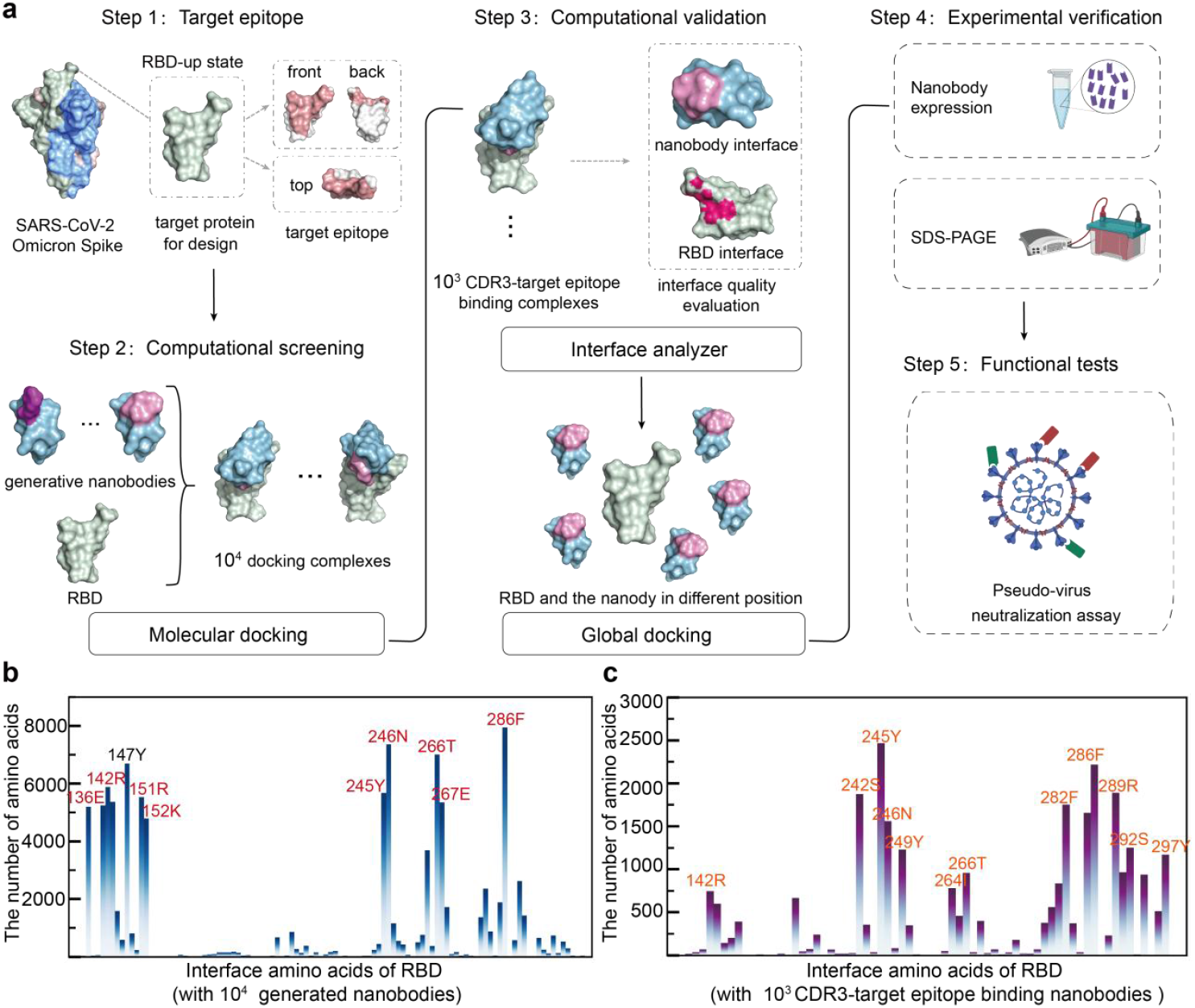
Recognition and validation of CDR3-epitope binding nanobodies. (a) Screening of site-specific nanobodies. Step I: Determination of the target epitope. The target protein is represented as a surface (light green), with the epitope represented in purple.Step II: Computational screening.The 10^4^ generated nanobodies were docked with the SARS-CoV-2 Omicron RBD using HDOCK software, and after obtaining 10^4^ docked complexes, we screened the complexes that bound with the target epitope and CDR3 (10^3^). This was done in order to identify those complexes that were most likely to bind with the target epitope and CDR3. Step 3, Computational validation. The interface of the 10^3^ selected complexes was scored using the Rosetta interface analyzer module. Complex structures with a high score in Rosetta interface analyzer module were then docked with the RBD using Rosetta global docking, and their binding energy landscapes were analyzed. Step 4, Experimental verification. The computationally evaluated nanobodies will be used for experimental verification, including the expression and purification of the nanobodies. Step 5, Functional tests. Successfully expressed and purified nanobodies will continue to be used in pseudovirus (Omicron) Neutralization to verify their activity. (b) Docking results of RBD with all nanobodies. The statistical results of amino acids at the interface on the RBD following the docking of 10^4^ nanobodies with the RBD. The amino acids labelled in red are the target epitope amino acids. (c) Docking results of RBD with 10^3^ nanobodies. The statistical results of amino acids at the interface on the RBD in 10^3^ complexes with CDR3-target epitope as the binding interface. All interface amino acids belong to the target epitope amino acids.

After defining the target binding sites, we applied a structured workflow to screen the library. As shown in Fig. 4a, we first used HDOCK (*28*) to dock each of the 10^4^ nanobodies from the library with the RBD. We then selected the top-ranking configuration from each of the 9,984 docking results based on docking scores and calculated amino acids involved in interactions between the nanobodies and the RBD within a 4 Å radius. We found that, apart from 147Y, all high-frequency RBD residues involved in nanobody binding were located within our target epitope (Fig. 4b), confirming that these sites on the “up” RBD serve as the primary binding locations for nanobodies (Fig. S5a). To further examine interactions involving the CDR3 and the RBD target epitope, we calculated the solvent-accessible surface area (SASA, Materials and Methods) and identified 2,783 complexes for which CDR3 and the RBD epitope formed the binding interface. We then analyzed these complexes and found that RBD residues at the interface were within the target epitope, with 242S, 245Y, 246N, 249Y, 282F, 286F, 289R, 292S, and 297Y being high-frequency residues involved in nanobody interactions (Fig. S5b).

Following this, we ranked the 10^3^ complexes based on HDOCK docking scores and retained the top 16 (docking score < -700) for further evaluation. We used the Rosetta interface analyzer (*29*) to assess the quality of these 16 complexes. Interface evaluation (Table S2) indicated that 12 nanobodies met the quality criteria (dG_separated/dSASA x 100 < -1.5, packstat > 0.65). (*29*) To validate these 12 docking results, we conducted global docking for each nanobody, using the initial configuration obtained from HDOCK, and performed 2,000 docking simulations using Rosetta global docking. (*30*) As shown in Fig. S6, low-interface-score conformations had RMSD values near zero, clustering in the lower left of the plot, which suggests that the lowest-energy conformations aligned with the initial poses, indicating consistency between the globally docked complex structures and the HDOCK results. Among these complexes, several nanobody-RBD conformations exhibited multiple RMSD values close to zero at the lowest energy. Ten nanobodies had docking complexes more similar to the initial poses and showed lower energy values, whereas Nb4759 and Nb4114 each had only one conformation at the lowest energy. We inferred that multiple low-RMSD conformations at the lowest energy might imply a stronger potential interaction between the nanobody and RBD at these sites. Consequently, we selected these ten nanobodies for subsequent experimental validation.

### Pseudovirus Neutralization Validation

To validate the computational findings, functional experiments were conducted on the ten candidate nanobodies. Initially, the expression of these ten nanobodies was induced in *Escherichia coli* Rosetta (DE3), followed by purification using Ni-NTA, successfully resulting in the acquisition of all ten nanobodies (Fig. 5a).

**Figure 5.**
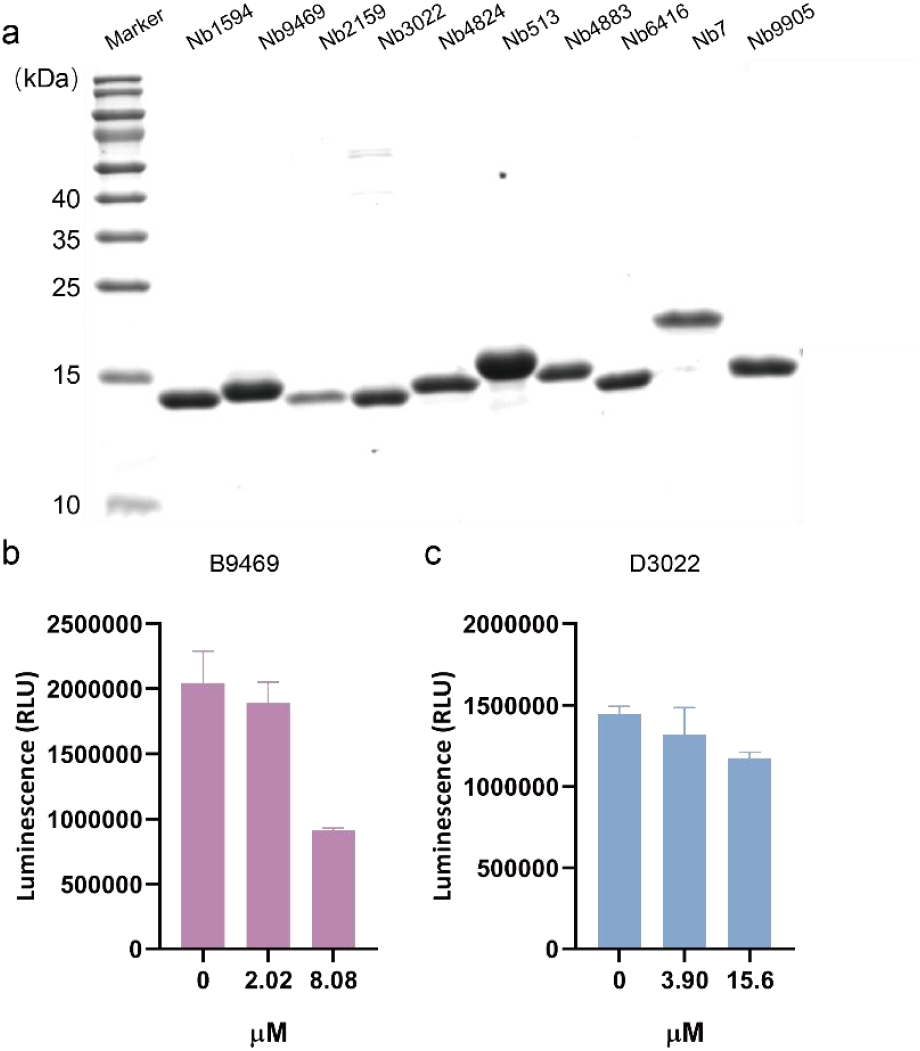
Functional tests for nanobodies. (a) Reducing SDS-PAGE of 10 nanobodies purified from E. coli Rosetta (DE3) cells. (b-c) Antiviral neutralization against the SARS-CoV-2 pseudoviruses (Omicron) by 2 nanobodies, respectively (biological replicate n = 3).

Next, we assessed the neutralization activity of these ten nanobodies against SARS-CoV-2 pseudovirus (Omicron BA.1). As shown in Fig. 5b-c, the luminescence signal gradually decreased with increasing concentrations of Nb9469 and Nb3022. At a concentration of 8.08 μM, the luminescence signal of Nb9469 was reduced by half, to 911,595 RLU (Fig. 5b). Similarly, at a concentration of 15.6 μM, the luminescence signal of Nb3022 decreased to 1,174,755 RLU (Fig. 5c). This result confirmed that among the ten nanobodies (Fig. 5b-c and Fig .S7b-d), Nb9469 and Nb3022 indeed exhibited neutralization potential against Omicron.

Through these functional experiments, we successfully validated that by modifying only the CDR3 region of the nanobody, the entire nanobody could acquire neutralization potential against the Omicron variant. However, it is important to note that while the neutralization was detectable, the overall neutralization potency was relatively weak, as indicated by the high IC50 values of both nanobodies. Given the limited neutralization ability, further experiments such as binding affinity studies (e.g., K_D_ measurements) were not pursued, which constrained our ability to perform direct comparisons with other nanobodies.

### Epitope Mapping and Interaction Analysis

To further investigate the molecular basis of the neutralization potential of Nb9469 and Nb3022, we performed an interface analysis between these two nanobodies and the Omicron RBD, using the docking results from the screening process. As shown in Fig. 6a-b, both nanobodies bind to distinct regions of the Omicron RBD.

**Figure 6.**
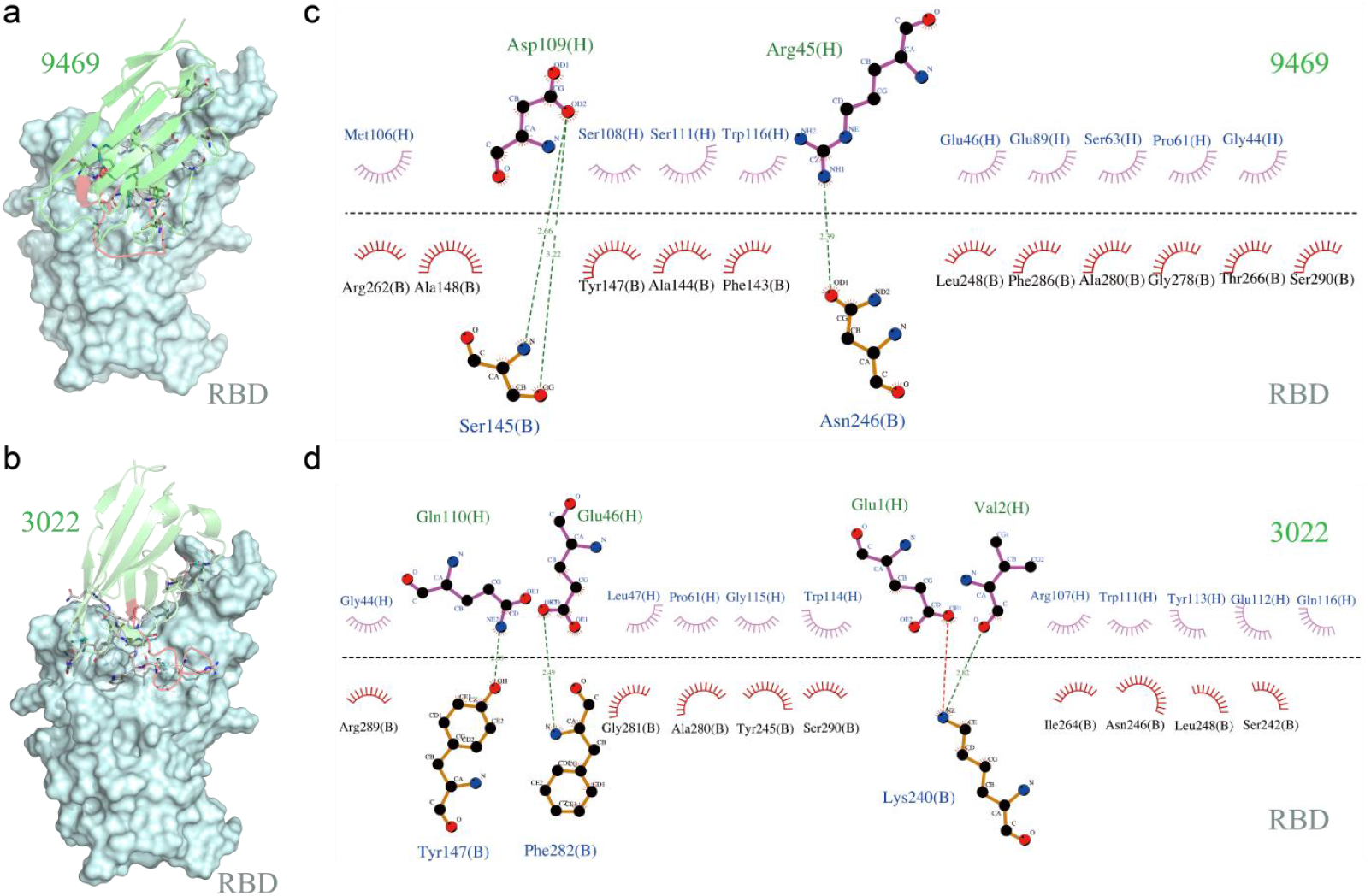
Interaction interfaces between neutralizing nanobodies and the target protein. (a) Overall structure of the Nb9469-RBD complex formed by docking nanobody 9469 with the Omicron RBD protein. Nb9469 (cyan) and its CDR3 (red) are shown in cartoon representation, while the RBD (light blue) is displayed using a surface representation. Residues involved in interactions are shown as stick models. (b) Overall structure of the Nb3022-RBD complex formed by docking nanobody 3022 with the Omicron RBD protein. Nb3022 (cyan) and its CDR3 (red) are shown in cartoon representation, while the RBD (light blue) is displayed using a surface representation. Residues involved in interactions are shown as stick models. (c-d) Detailed interactions between the CDR3s of 9469 and 3022 and the Omicron RBD. Interacting residue pairs are displayed as stick models, with hydrogen bonds represented by green dashed lines.

Nanobody Nb9469 forms interactions primarily with residues located in the central region of the Omicron RBD. Detailed interface analysis using LigPlot+ (*31*) reveals key interactions between nanobody Nb9469 and the RBD (Fig. 6c). Nanobody Nb9469 primarily interacts with the RBD through hydrogen bonds and hydrophobic interactions. Key hydrogen bonds are formed between Asp109(H) and Ser145(B), as well as Arg45(H) and Asn246(B) (Fig. 6c). Additionally, Ser108(H) and Trp116(H) form hydrogen bonds with Ala148(B) and Phe143(B), respectively. These interactions are complemented by numerous hydrophobic contacts, particularly with residues Leu248(B), Ala280(B), and Gly278(B). Together, these interactions stabilize the binding of Nb9469 to the RBD, potentially contributing to its neutralization ability.

Nanobody Nb3022, on the other hand, engages a different set of residues on the RBD. Its binding is mediated by hydrogen bonds formed between Gln110(H) and Glu46(H), as well as Glu1(H) and Lys240(B) (Fig. 6d). Additional interactions are observed between Val2(H) and Ser290(B), stabilizing the interface. Hydrophobic interactions, particularly with Phe282(B) and Tyr245(B), further strengthen the binding between Nb3022 and the RBD. The difference in binding sites and interaction profiles between Nb9469 and Nb3022 indicates that these two nanobodies may neutralize the Omicron variant through distinct mechanisms.

These structural analyses provide molecular insights into how selective modification of the CDR3 region can yield nanobodies with neutralizing potential.

## Discussion

In this study, we present a de novo nanobody design method based on GAN, offering a scalable and efficient approach to nanobody discovery. By leveraging deep learning, we generated a large nanobody library and identified candidates with promising target specificity. Using our AiCDR model, we generated approximately 10^4^ natural-like CDR3 sequences, which we grafted onto nanobody scaffolds to construct a structural library containing roughly 10^4^ nanobodies. Employing docking techniques, we further identified ten nanobodies with favorable binding interfaces to the RBD of the SARS-CoV-2 Omicron variant. Functional assays confirmed that two of these nanobodies demonstrated some degree of neutralization activity against the Omicron variant in vitro.

This method offers advantages over traditional nanobody discovery methods. Through the AiCDR model, we have achieved high efficiency, low-cost library construction that avoids the ethical and time constraints associated with animal immunization. Although GANs have been increasingly applied to biological sequence generation, AiCDR is the first model specifically tailored for nanobody CDR3 sequence generation. The inclusion of two Guider modules in AiCDR enhances its capacity to filter out non-functional sequences, producing sequences that are both natural-like and highly diverse. This high-quality sequence diversity enhances the potential for targeting a range of antigenic epitopes.

Furthermore, the structural library constructed here not only provides nanobody sequence data but also comprehensive structural information, facilitating future nanobody screening against other target proteins. This library serves as a valuable tool for researchers targeting other proteins, such as respiratory syncytial virus (RSV), thereby broadening the scope of nanobody screening. Incorporating computational screening further reduces the complexity and randomness inherent in experimental screening methods.

While we identified two neutralizing nanobodies among ten candidates, their neutralization capacity was relatively weak. Additional modifications to the CDR1 and CDR2 regions could further enhance their antiviral efficacy. Both regions contribute to antigen binding, and optimizing CDR1, CDR2, and CDR3 together may improve binding affinity and neutralizing capacity against the Omicron Spike protein. This multi-region optimization approach could significantly enhance the efficacy of nanobodies and provide a rapid response to evolving SARS-CoV-2 variants.

The AiCDR model also holds potential for broader applications. Beyond CDR3 sequence generation, adjustments to the training data could allow AiCDR to generate CDR1, CDR2, or even complete nanobody sequences. This model could further extend to other biological sequences, such as peptides. In addition, the screening process we developed can be adapted for use with any target protein of known structure, offering a versatile platform for antibody design against diverse pathogens.

Recent advances in diffusion (*32*) models have also shown great promise in biological applications, including those in AlphaFold3, (*33*) where diffusion-based models have become powerful tools for molecular generation. Applications have emerged in generating 3D molecular structures from protein pocket features, (*34*) molecular docking tools like DiffDock, (*35*) and antibody design. Moving forward, we plan to integrate such advanced generative models to enhance sample quality and diversity, along with implementing these deep learning tools in subsequent screening processes to streamline and enhance our workflow.

In conclusion, this study proposes a GAN-based method for the de novo design of nanobodies. This approach generated natural-like CDR3 sequences and identified nanobodies capable of neutralizing the Omicron Spike protein. It introduces a new method for constructing antibody libraries and offers a foundational resource for computational nanobody design, which may facilitate the discovery of target-specific nanobodies. Additionally, our screening approach provides an efficient means to identify nanobodies with neutralizing potential, representing a promising step forward in antibody development.

## Materials and methods

### Screening of Complexes with CDR-Target as Binding Interface

We used HDOCK to dock 9984 nanobodies with Omicron RBD. Each nanobody formed multiple conformations upon docking with RBD. We selected the conformation with the lowest HDOCK Docking score as the optimal conformation for each nanobody-RBD interaction, resulting in 9984 optimal docking conformations. Subsequently, we computed the average area difference (Δ Scdr3) for the nanobody CDR3 before and after binding to RBD. This step involved calculating the accessible surface area of CDR3 before and after binding to RBD using Biopython’s Bio.PDB.SASA module, and dividing it by the length of CDR3. Conformations with ΔScdr3=0 were considered non-CDR3 binding conformations. Next, we performed a similar calculation for Omicron before and after binding to the nanobodies to determine the area difference (ΔStarget). Conformations with ΔStarget = 0 were considered non-target binding conformations. We retained complexes with ΔScdr3 > 0 and ΔStarget > 0 as CDR-target binding interface complexes.

### Rosetta Interface Analyzer for Complex Interface Scoring

We have prepared a blueprint file to specify the required computations, such as whether to run the packstat calculation in Rosetta. We set parameters such as ‘-compute_packstat’, ‘-tracer_data_print’, ‘-pack_input’, ‘-pack_separated’, and ‘-add_regular_scores_to_scorefile’ to ‘true’, while leaving other parameters at their default or recommended values. This blueprint is utilized to evaluate the interface quality of each nanobody-RBD complex, generating a score file. Through this score file, we can assess whether the interface of the complex is satisfactory (with criteria such as dG_separated/dSASAx100 < -1.5 and packstat > 0.65).

### Validation of docking consistency using Rosetta Global Docking

For the 12 nanobody-RBD complexes that exhibited good interfacial scores, we utilized their optimal conformations from HDOCK docking as the initial conformations. Employing Rosetta for global docking, we conducted five parallel docking sessions, each consisting of 1000 docking attempts, totaling 5000 attempts. In our protocol, we specified the parameters as follows: ‘-nstruct’ set to 1000, ‘-partners A_B’ (where A represents the nanobody and B the RBD), ‘-dock_pert 3 8’, ‘-spin -randomize1-ex1-ex2aro’. Upon completion of these global dockings, we delineated the binding energy landscape for each complex using the ‘I_sc’ and ‘Irms’ parameters from the output score file.

Based on the dot plot of the binding energy landscape, we checked 2000 interface scores and RMSDs in comparison to the initial conformation for each complex. This analysis allowed us to select the best or several superior complexes for subsequent experimental validation.

### Protein expression and purification

The *Escherichia coli* (E. coli) bacterium was utilized to express the candidate nanobodies. The codon-optimized gene encoding the designed proteins was cloned into a pET-26b (+) vector, which included an N-terminal 8 × His-tag and a TEV cleavage site. These vectors were then introduced into E. coli Rosetta (DE3) cells. The cells were cultured in shake flasks containing LB broth with kanamycin and chloramphenicol at 37°C until the optical density at 600 nm (OD600) exceeded 1. Isopropyl-β-D-thiogalactoside (IPTG) was subsequently added, and the flasks were incubated at 25°C for 20 hours. Periplasmic proteins were extracted using an osmotic shock method. The resulting supernatant was filtered through a 0.45 µm syringe filter, and the target proteins were purified using a gravity-flow column containing 2 mL of Ni-NTA resin. The column was initially equilibrated with buffer A (pH 7.4, 1×PBS, 20 mM imidazole), followed by the addition of the filtered supernatant. Elution was performed using buffer A and buffer B (pH 7.4, 1×PBS, 300 mM imidazole), and the protein was finally eluted with buffer B. The purified proteins were analyzed by sodium dodecyl sulfate-polyacrylamide gel electrophoresis (SDS-PAGE). To remove the His-tag, the purified proteins were desalted using a HiTrap desalting column and incubated overnight at 4°C with TEV protease. The tag-free protein was then obtained using a Ni-NTA gravity-flow column.

### Neutralization assay with SARS-CoV-2 spike BA. 1 pseudotyped viruses

The pseudovirus neutralization assays did by ACROBiosystems Inc. The SARS-CoV-2 Spike pseudoviruses carrying firefly luciferase reporter gene were generated with vesicular stomatitis virus (VSV) pseudotyping system according to a published protocol. (*36*) Briefly, plasmids encoding propotype or a variant SARS-CoV-2 spike protein BA. 1, was transfected into HEK 293T cells together with VSVΔG pseudovirus, respectively. Next day, viral supernatants were harvested at 24 h post-infection and then centrifuged and filtered to remove cell debris and stored at −80°C. Virus titers were calculated by testing the median tissue culture infective dose with a human ACE2 overexpression cell line. (*36*) For neutralization assay, the pseudoviruses were diluted to approximately 10^4^ TCID_50_/mL with the cell culture medium. Then, 75 μL protein sample was mixed with 25 μL SARS-CoV-2 Spike pseudovirus dilution in a 96-well white flat bottom plate, and incubated at 37 °C for 1 hour, followed by the addition of 100 μL of human ACE2 overexpression HEK 293 cell suspension (5×10^4^ cells/well) to the mixtures. After incubation at 37°C, 5% CO_2_ (v/v) for 24 hours, the luminescence value was detected with the aid of luminescence detection reagent and a luminescence meter (PerkinElmer, Cat. No. HH34000000). The inhibition ratio was calculated with the following formula:

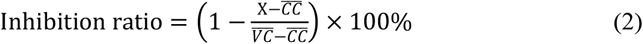

X: the luminescence value (RLU) of a certain well;

CC, cell control, only cells are added;

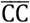, the mean value of cell control group;

VC, virus control, only cells and pseudovirus are added;

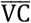, the mean value of virus control group.

Each assay was done in triplicate. Data were fitted using non-linear regression, and the ID_50_ values were calculated using a four-parameter regression equation in GraphPad Prism.

## Supplementary information

The online version contains supplementary material available at …

## Contributions

Q.H. supervised the project. L.H. and Q.H. conceived the project; L.H. performed computational analysis; T.T., L.H. Y.X., Q.Q., X.J. performed the experiments; L.H. and Q.H. wrote the manuscript.

## Availability of data and materials

All data generated or analyzed during this study have been included in the article and the Additional information.

## Declarations

### Ethics approval and consent to participate

Not applicable.

