## Supplementation for "Deep generative design of neutralizing nanobodies against SARS-CoV-2 variants"

Deep learning-driven CDR3 design for neutralizing

nanobodies against SARS-CoV-2

***List of Supplementary Materials***

Figure S1. The training process of Guider1 and Guider2.

Figure S2. The training perfomance of different combination of α、β、γ.

Figure S3. The structures of four nanobodies predicted by Alphafold2.

Figure S4. The Omicron S protein in the 1 RBD-up, 2 RBD-down conformation.

Figure S5. Binding Epitope Map of Nanobody and RBD Docking.

Figure S6. Binding free energy landscape of candidate nanobodies.

Figure S7. Functional test of Caplacizumab and eight nanobodies with negative results.

Table S1. The combinations of α, β, and γ values.

Table S2. The cluster of generated sequences and training dataset.

Table S3. The cluster group of generated sequences and training dataset.

Table S4. The good interface analyzer results of 12 nanobodies respectivily.

Table S5 . The amino acid sequences of the designed nanobodies for experimental tests.

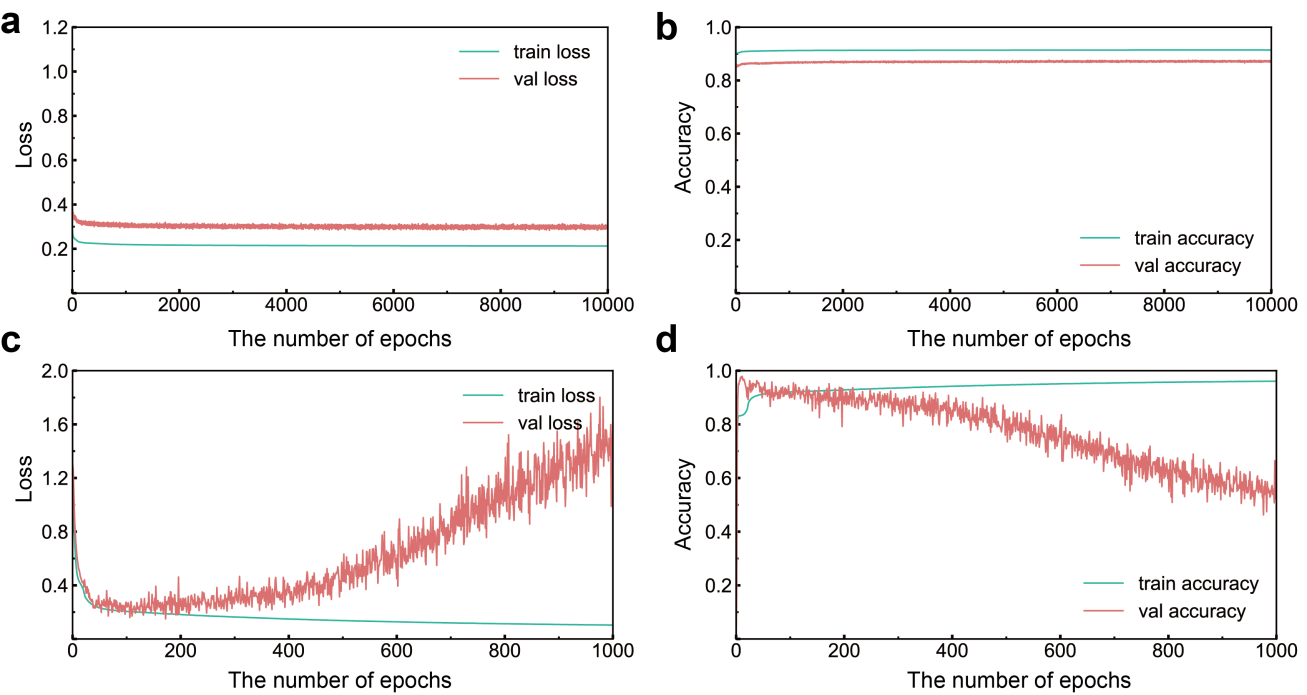

**Figure S1.** The training process of Guider1 and Guider2. **(a)** The loss of Guilder1 during the training process. **(b)**  The accuarcy of Guilder1 during the training process. **(c)** The loss of Guilder2 during the training process. **(d)** The accuracy of Guilder2 during the training process.

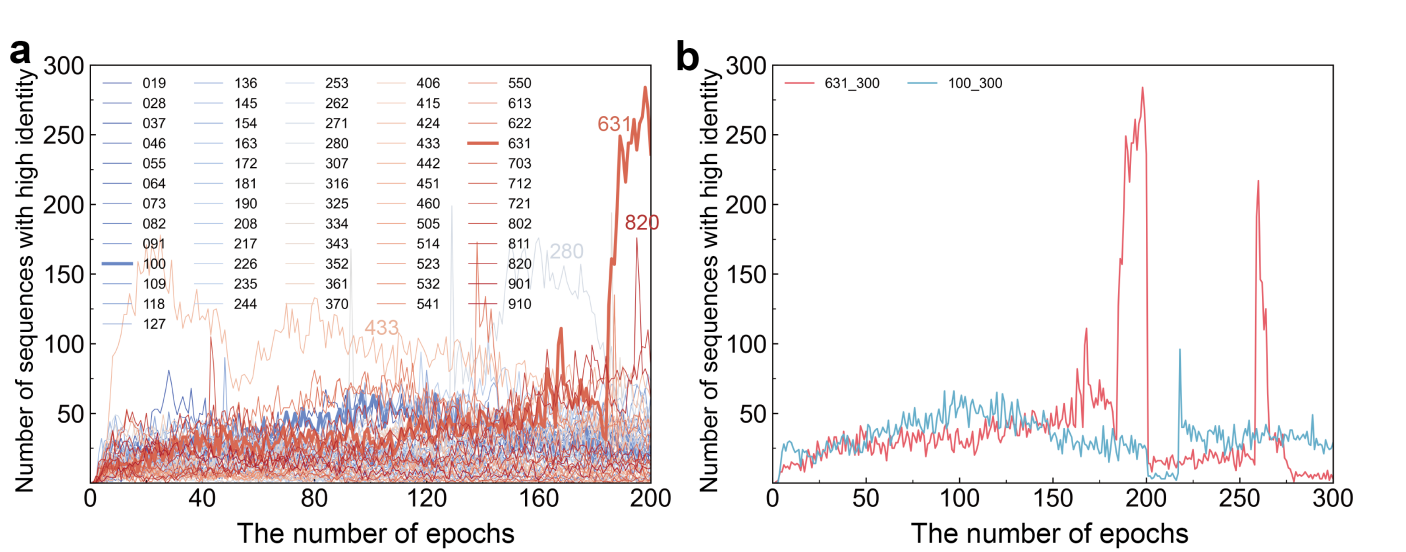

**Figure S2.** Training effect of AiCDR with different α , β, and γ combinations. (a) Comparison of the number of similar sequences generated by different networks. The figure represents the number of sequences similar to native sequences generated by different networks during the training process. Different colors represent AiCDR networks with different α, β, γ combinations, and there are 61 groups in total. (b) Comparison of the number of similar sequences generated by 631 and 100 networks. The number of similar sequences generated after running 631 and 100 networks for another epochs (300 epochs in total).

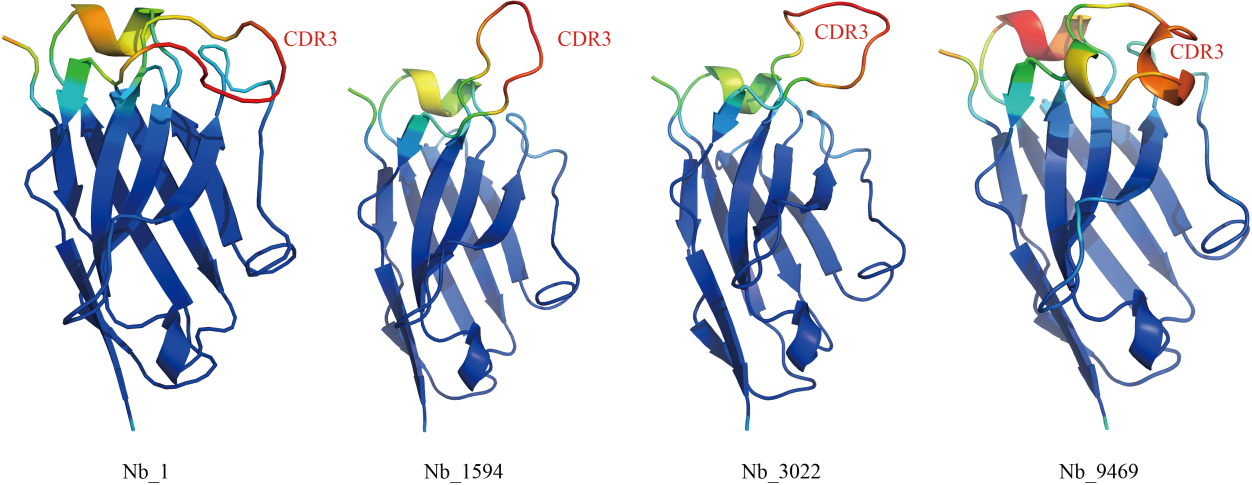

**Figure S3.** The structures of four nanobodies predicted by Alphafold2 shown in cartoon representation and color-coded from blue to red based on pLDDT scores, ranging from high to low confidence. The nanobody framework regions display high pLDDT scores, while the CDR regions show lower pLDDT values, with the CDR3 region having the lowest scores.

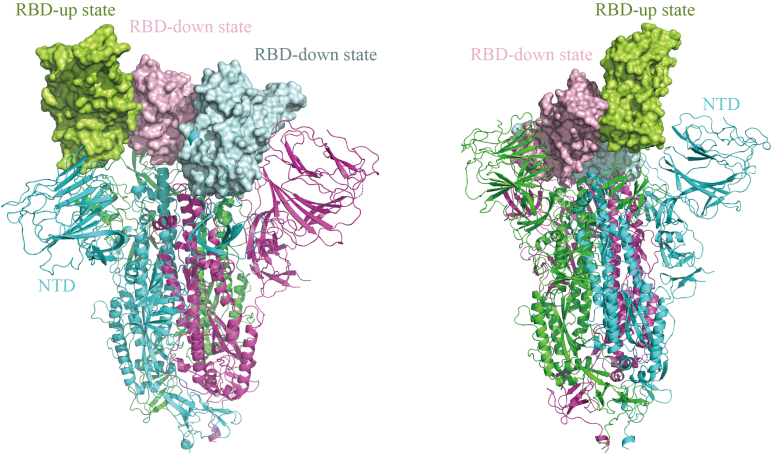

**Figure S4.** The Omicron S protein in the 1 RBD-up, 2 RBD-down conformation (PDB: 7TGW), with RBD regions shown in distinct colors on the surface.

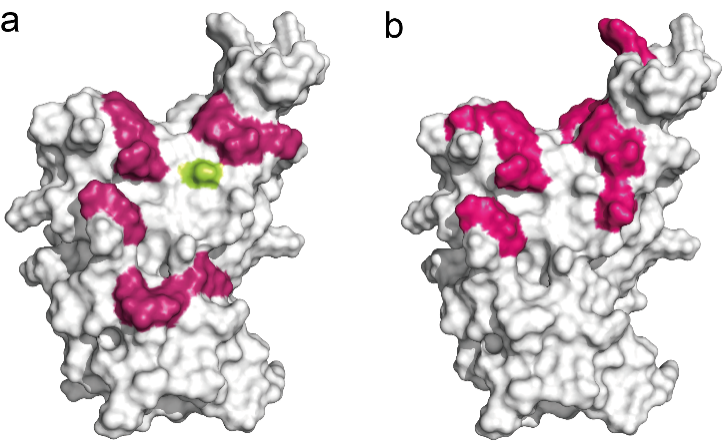

**Figure S5.** Binding Epitope Map of Nanobody and RBD Docking. (a) Binding epitope representation on RBD when docked with 104 nanobodies. Red-labeled amino acids indicate target binding sites, while yellow-labeled amino acids denote non-target binding sites. (b) Binding epitope representation on RBD when docked with 10^3^ nanobodies. Red-labeled amino acids indicate target binding sites.

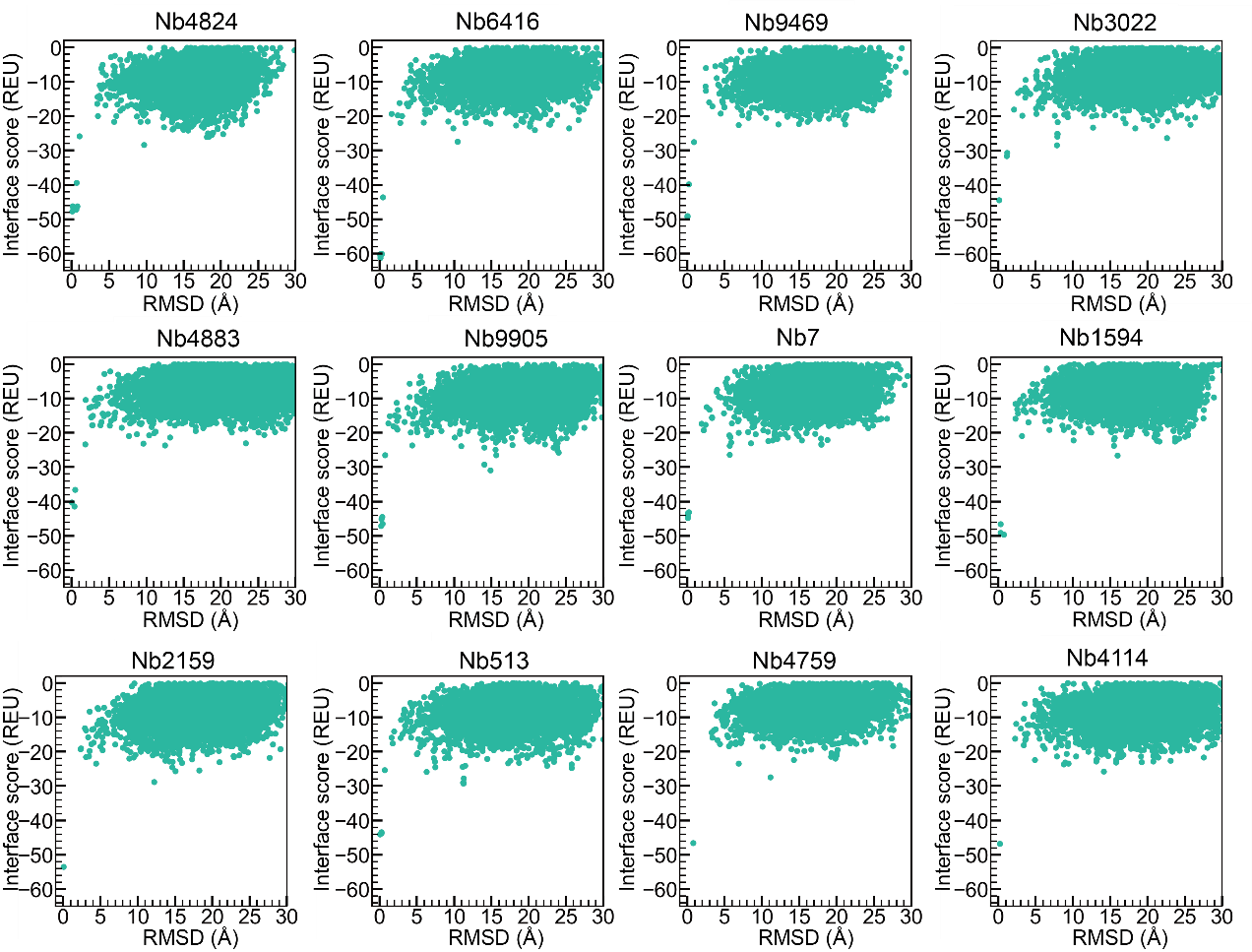

**Figure S6.** Binding free energy landscape of candidate nanobodies. The binding free energy landscape obtained after docking SARS-CoV-2 Omicron with 12 nanoantibodies using Rosetta global docking with RMSD in the horizontal coordinate and Interface score (REU) in the vertical coordinate indicates that during global docking, the structure of the complexes with respect to the initial conformational RMSD of the complex structure relative to the initial conformation during global docking and its interface score at the corresponding position.

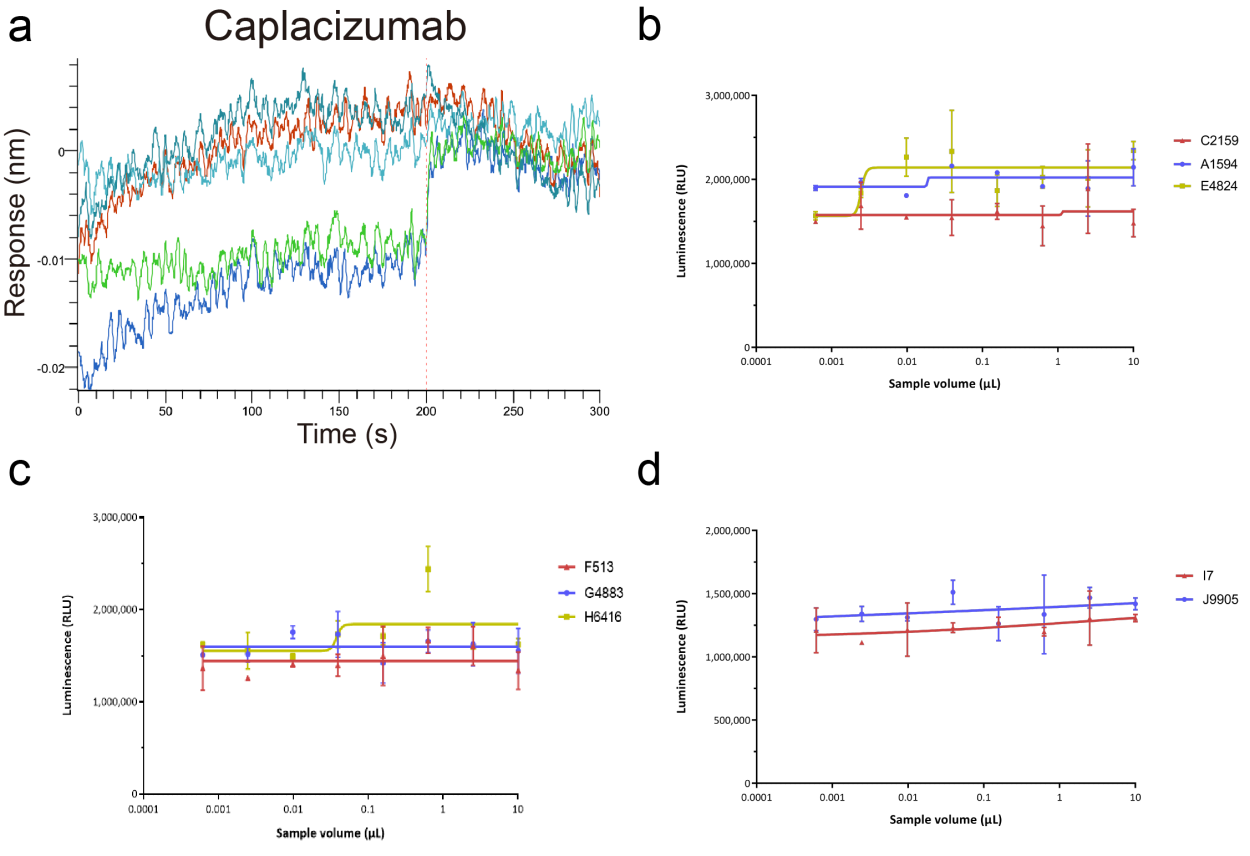

**Figure S7.** Functional test of Caplacizumab and eight nanobodies with negative results. (a) BLI measurements for Caplacizumab with Omicron RBD. (b-d) Antiviral neutralization against the SARS-CoV-2 pseudoviruses (Omicron) by 8 nanobodies, respectively (biological replicate n = 3).

**Table S1.** The combinations of α, β, and γ values.

|  | Combinations (61 in total) |
| --- | --- |
| **(**α, β, γ**)** | (0, 0.1, 0.9), (0, 0.2, 0.8), (0, 0.3, 0.7), (0, 0.4, 0.6), (0, 0.5, 0.5), (0, 0.6, 0.4), (0, 0.7, 0.3), (0, 0.8, 0.2), (0, 0.9, 0.1), (0.1, 0, 0.9), (0.1, 0.1, 0.8), (0.1, 0.2, 0.7), (0.1, 0.3, 0.6), (0.1, 0.4, 0.5), (0.1, 0.5, 0.4), (0.1, 0.6, 0.3), (0.1, 0.7, 0.2), (0.1, 0.8, 0.1), (0.1, 0.9, 0), (0.2, 0, 0.8), (0.2, 0.1, 0.7), (0.2, 0.2, 0.6), (0.2, 0.3, 0.5), (0.2, 0.4, 0.4), (0.2, 0.5, 0.3), (0.2, 0.6, 0.2), (0.2, 0.7, 0.1), (0.2, 0.8, 0), (0.3, 0, 0.7), (0.3, 0.1, 0.6), (0.3, 0.2, 0.5), (0.3, 0.3, 0.4), (0.3, 0.4, 0.3), (0.3, 0.5, 0.2), (0.3, 0.6, 0.1), (0.3, 0.7, 0), (0.4, 0, 0.6), (0.4, 0.1, 0.5), (0.4, 0.2, 0.4), (0.4, 0.3, 0.3), (0.4, 0.4, 0.2), (0.4, 0.5, 0.1), (0.4, 0.6, 0), (0.5, 0, 0.5), (0.5, 0.1, 0.4), (0.5, 0.2, 0.3), (0.5, 0.3, 0.2), (0.5, 0.4, 0.1), (0.5, 0.5, 0), (0.6, 0.1, 0.3), (0.6, 0.2, 0.2), (0.6, 0.3, 0.1), (0.7, 0, 0.3), (0.7, 0.1, 0.2), (0.7, 0.2, 0.1), (0.8, 0, 0.2), (0.8, 0.1, 0.1), (0.8, 0.2, 0), (0.9, 0, 0.1), (0.9, 0.1, 0), (1, 0, 0) |

**Table S2.** The cluster of generated sequences and training dataset.

| Training dataset (1,702,270 sequences) | Generated sequences (9,984 sequences) |
| --- | --- |
| 29,066 clusters | 1,308 clusters |

Note: using cd-hit (identity=0.5)

**Table S3.** The cluster group of generated sequences and training dataset.

| Data set | Cluster group |
| --- | --- |
| Training dataset  (1,702,270 sequences) | 1 ~ 4， 5 ~ 10， 11 ~ 20， 21 ~ 40， 41 ~80，81~ 160，161 ~ 320， 321 ~ 700， 701 ~ 1800， 1801 ~ 12727 |
| Generated sequences  (9,984 sequences) | 1 ~ 4， 5 ~ 10， 11 ~ 20， 21 ~ 40， 41 ~80，81~ 160，161 ~ 320， 321 ~ 700 |

Note: Clusters containing 1 to 4 sequences are grouped into a single cluster group labeled "1 ~ 4."

**Table S4.** The interface analyzer results of 12 nanobodies respectivily (ranking by HDOCK energy).

| Nb | HDOCK | CDR3 | total | dG | dG/dSASA x100 | dSASA | unsat | Hbonds | nres_int | Packstat |
| --- | --- | --- | --- | --- | --- | --- | --- | --- | --- | --- |
|  | energy | SASA | score | separated | (< -1.5) |  | Hbonds | int |  | (> 0.65) |
| 4321 | -766 | 39.5 | -862 | -46.1 | -2.07 | 2234 | 24 | 9 | 60 | 0.629 |
| 1594 | -737 | 27.0 | -888 | -50.6 | **-2.47** | 2050 | 11 | 14 | 44 | **0.667** |
| 9469 | -730 | 24.7 | -884 | -45.2 | -2.45 | 1843 | 16 | 7 | 50 | 0.647 |
| 4908 | -723 | 24.9 | -905 | -59.9 | -2.47 | 2424 | 19 | 17 | 51 | 0.623 |
| 2159 | -719 | 15.2 | -867 | -50.2 | **-2.57** | 1951 | 10 | 9 | 49 | **0.693** |
| 6795 | -716 | 18.6 | -914 | -32.8 | -1.95 | 1682 | 10 | 8 | 44 | 0.644 |
| 3022 | -715 | 23.9 | -887 | -41.5 | **-2.12** | 1956 | 15 | 8 | 48 | **0.672** |
| 4824 | -715 | 28.3 | -907 | -44.5 | **-2.45** | 1821 | 17 | 10 | 43 | **0.667** |
| 4873 | -711 | 24.1 | -883 | -52.0 | **-2.43** | 2146 | 17 | 12 | 59 | **0.685** |
| 4114 | -709 | 30.9 | -884 | -40.3 | **-2.24** | 1803 | 15 | 11 | 37 | **0.695** |
| 513 | -708 | 20.6 | -887 | -41.5 | **-2.40** | 1729 | 13 | 13 | 40 | **0.657** |
| 792 | -706 | 24.7 | -861 | -41.5 | **-2.09** | 1983 | 16 | 8 | 52 | **0.673** |
| 4883 | -705 | 20.8 | -885 | -43.9 | **-2.09** | 2097 | 22 | 8 | 53 | **0.656** |
| 4759 | -704 | 27.1 | -885 | -50.7 | **-2.28** | 2228 | 14 | 13 | 56 | **0.651** |
| 6416 | -703 | 26.2 | -911 | -57.0 | **-2.92** | 1954 | 7 | 12 | 47 | **0.684** |
| 7 | -702 | 26.2 | -889 | -42.3 | **-2.13** | 1986 | 18 | 10 | 49 | **0.710** |

**Table S5.** The amino acid sequences of the designed nanobodies for experimental tests.

| Name | Amino acid sequence |
| --- | --- |
| 1594 | EVQLVESGGGLVQPGGSLRLSCAASGRTFSYNPMGWFRQAPGKGRELVAAISRTGGSTYYPDSVEGRFTISRDNAKRMVYLQMNSLRAEDTAVYYCAALYARSGGVEYQYDYWGQGTQVTVSS |
| 2159 | EVQLVESGGGLVQPGGSLRLSCAASGRTFSYNPMGWFRQAPGKGRELVAAISRTGGSTYYPDSVEGRFTISRDNAKRMVYLQMNSLRAEDTAVYYCAADGRTVHTRWGLLEVEFGYWGQGTQVTVSS |
| 6469 | EVQLVESGGGLVQPGGSLRLSCAASGRTFSYNPMGWFRQAPGKGRELVAAISRTGGSTYYPDSVEGRFTISRDNAKRMVYLQMNSLRAEDTAVYYCAASPQPLPAMYSDLSVYEYWGQGTQVTVSS |
| 3022 | EVQLVESGGGLVQPGGSLRLSCAASGRTFSYNPMGWFRQAPGKGRELVAAISRTGGSTYYPDSVEGRFTISRDNAKRMVYLQMNSLRAEDTAVYYCSAADFRLVVGRTPQWEYWGQGTQVTVSS |
| 4824 | EVQLVESGGGLVQPGGSLRLSCAASGRTFSYNPMGWFRQAPGKGRELVAAISRTGGSTYYPDSVEGRFTISRDNAKRMVYLQMNSLRAEDTAVYYCAADRRPCSGLRHGEYDYWGQGTQVTVSS |
| 513 | EVQLVESGGGLVQPGGSLRLSCAASGRTFSYNPMGWFRQAPGKGRELVAAISRTGGSTYYPDSVEGRFTISRDNAKRMVYLQMNSLRAEDTAVYYCAVDDDEAEGLVRDGYWGQGTQVTVSS |
| 4883 | EVQLVESGGGLVQPGGSLRLSCAASGRTFSYNPMGWFRQAPGKGRELVAAISRTGGSTYYPDSVEGRFTISRDNAKRMVYLQMNSLRAEDTAVYYCAADLFCVYSDSQWYGDYDYWGQGTQVTVSS |
| 6416 | EVQLVESGGGLVQPGGSLRLSCAASGRTFSYNPMGWFRQAPGKGRELVAAISRTGGSTYYPDSVEGRFTISRDNAKRMVYLQMNSLRAEDTAVYYCARGRAQLGDFGPGYPDPWGQGTQVTVSS |
| 7 | EVQLVESGGGLVQPGGSLRLSCAASGRTFSYNPMGWFRQAPGKGRELVAAISRTGGSTYYPDSVEGRFTISRDNAKRMVYLQMNSLRAEDTAVYYCAADEEHWCGDSWSGEYDYWGQGTQVTVSS |
| 9905 | EVQLVESGGGLVQPGGSLRLSCAASGRTFSYNPMGWFRQAPGKGRELVAAISRTGGSTYYPDSVEGRFTISRDNAKRMVYLQMNSLRAEDTAVYYCAAAAKPWTFFAGHWGQGTQVTVSS |
